# RNA sequencing of Stentor cell fragments reveals key processes underlying cellular regeneration

**DOI:** 10.1101/232363

**Authors:** Henning Onsbring Gustafson, Mahwash Jamy, Thijs J. G. Ettema

## Abstract

While ciliates of the genus *Stentor* are known for their ability to regenerate when their cells are damaged or even fragmented, the physical and molecular mechanisms underlying this process are poorly understood. To identify genes involved in the regenerative capability of *Stentor* cells, RNA sequencing of individual *Stentor polymorphus* cell fragments was performed. After splitting a cell over the anterior-posterior axis, the posterior fragment has to regenerate the oral apparatus, while the anterior part needs to regenerate the hold fast. Altogether, differential expression analysis of both posterior and anterior *S. polymorphus* cell fragments for four different post-split time points revealed over 10,000 up-regulated genes throughout the regeneration process. Among these, genes involved in cell signaling, microtubule-based movement and cell cycle regulation seemed to be particularly important during cellular regeneration. We identified roughly nine times as many up-regulated genes in regenerating *S. polymorphus* posterior fragments as compared to anterior fragments, indicating that regeneration of the anterior oral apparatus is a complex process that involves many genes. Our analyses identified several expanded groups of genes such as dual-specific tyrosine-(Y)-phosphorylation regulated kinases and MORN domain containing proteins that seemingly act as key-regulators of cellular regeneration. In agreement with earlier morphological and cell biological studies, our differential expression analyses indicate that cellular regeneration and vegetative division share many similarities.

## Results and discussion

The regenerative capabilities of the heterotrichous ciliate *Stentor* have been first documented over a century ago [1]. Since then, *Stentor* has become a model organism for studying cellular regeneration and repair, and, eventually, for developing new therapeutic approaches for curing human disease [2]. Yet, the molecular basis for *Stento?s* regenerative capability is largely unknown. Some pioneering studies employing microsurgery have revealed that regeneration and cell division share many similarities on nuclear level [3, 4]: Weisz observed macronuclear changes in posterior regenerating fragments, which he compared to dividing cells, while Guttes and Guttes found that the micronuclei in regenerating posterior fragments underwent mitosis. In this study, we employed RNA sequencing (RNAseq) of single *Stentor polymorphus* cells and cell fragments to gain insight into the molecular basis of cellular regeneration and to identify key determinants underlying this process. By employing RNAseq at the single cell or cell fragment level, we avoided having to culture and manipulate large numbers of *S. polymorphus* cells. It also allowed us to record any potential expression profile variations between individual cells and cell fragments (which were to be expected given the rather crude manual ‘microsurgeries’ that were performed to split individual cells).

Individual cells from a near-clonal *S. polymorphus* population were isolated from a pond in Uppsala (Sweden; see Methods for details), and subjected to microsurgeries during which cells were split over their axial plane, leaving an anterior cell fragment that contained the oral apparatus, and a posterior cell fragment containing the foothold. Observations of these regenerating *S. polymorphus* cell fragments indicated that the initial cell shape was regained after 6 hours. To study the regeneration process of anterior and posterior fragments at the transcriptional level, we employed an experimental setup in which RNAseq was performed for both fragments at 10 min, 90 min, 180 min and 300 min post split (Fig. 1a), in which each replicate was generated from a single regenerating *Stentor* fragment (Fig. 1b). The first time point, 10 min, was selected to characterize early expression after wound healing. We used 300 min as the last time point since *S. polymorphus* cells were observed to have regained the native shape after 360 min of regeneration. Additionally, two time points was used in between (90 and 180 min). The conditions corresponding to the four time points of regenerating anterior (A) replicates will be referred to as A10, A90, A180 and A300 while the conditions for the regenerating posterior (P) cell fragments will be referred to as P10, P90, P180, P300. As a reference, RNAseq was also performed to generate cDNA of non-regenerating *S. polymorphus* cells. For each of the conditions described above, four independent cDNA pools were generated (except the non-regenerating cells and P180, where only three replicates could be generated), followed by multiplexed library preparation and deep Illumina sequencing (see Methods for details).

**Figure 1.**
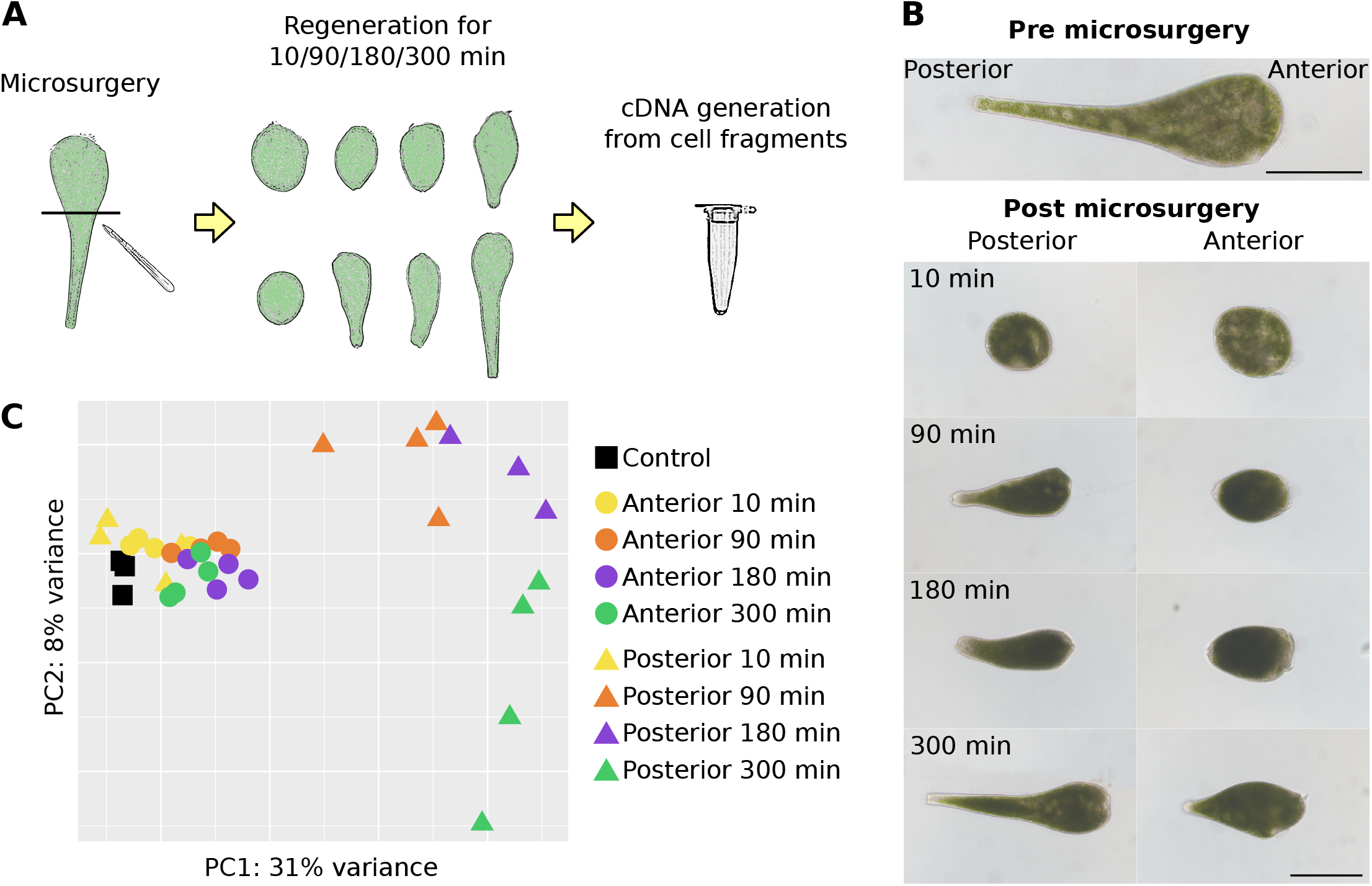
Overview of experimental design, cell morphology and clustering of differential expression data. a) cDNA was generated from single cell fragments. Eight different conditions were included in the experiment, corresponding to fragments regenerating until four different time points post split. Both anterior and posterior parts of the cell were analyzed. b) Photos of regenerating cell fragments at the different time points. Both scale bars correspond to 200 μm. c) PCA based on the expression level of all genes for each replicate included in the experiment. See also Table S2.

Paired Illumina sequence reads obtained from all libraries were co-assembled into a master transcriptome that contained 127,533 contigs. Next, to reduce the level of redundancy (e.g. due to splice variants, strain variation, miss-assembly, etc.), all contigs were clustered into 42,832 unique transcripts (see Methods for details). This number is roughly comparable to the 34,506 genes that were predicted in the recently published *S. coeruleus* genome [5]. We performed a principle component analysis of the gene expression levels across the samples to (i) assess whether replicates displayed similar regulation of genes and to (ii) determine whether samples with a different experimental background could be separated based on different expression levels (Fig. 1c). This analysis revealed that, overall, samples with the same experimental background tend to cluster in close proximity, indicating that expression variance between replicate experiments was within sustainable levels. Furthermore, we observed that regenerating posterior cell fragments (P90, P180 and P300) formed distinct clusters separated from all other conditions, and that, with the exception of the 10 min samples (P10, A10), the anterior and posterior replicates were all separated. These observations indicate that the obtained expression profiles reflect the experimental conditions biologically relevant clusters (also see Suppl. Information), encouraging us to functionally explore the transcriptomes of regenerating *S. polymorphus* cells in more detail.

Differential expression analysis of the transcriptome dataset revealed a total of 10,682 upregulated and 8,067 downregulated genes in all conditions when comparing to the non-regenerating *S. polymorphus* cells. In this study we have focused on the upregulated genes since these are likely to be involved in the repair of structures that *Stentor* is missing. In posterior fragments 6,433 genes were upregulated that were not upregulated in the anterior conditions, while only 720 upregulated genes were anterior-specific. Based on this observation, it is clear that regeneration of the oral apparatus represents a transcriptionally more complex process compared to regenerating the hold fast. Furthermore, our data indicates that the initial wound healing is a less transcriptionally complex process compared to cellular regeneration: only 9% (955 genes) and 10% (1117 genes) of all upregulated genes could be detected for A10 and P10, respectively. Among these, only 64 and 122 were specifically upregulated in A10 and P10, respectively. Each of the later stages of anterior fragment regeneration (A90, A180 and A300) seem to involve at least twice as many genes as compared to wound healing of the anterior fragment (A10; Fig. 2). For posterior regeneration, these numbers are even more pronounced, with at least five times as many genes being upregulated during each of the regenerating stages (P90, P180 and P300) as compared to wound healing (P10; Fig. 2).

**Figure 2.**
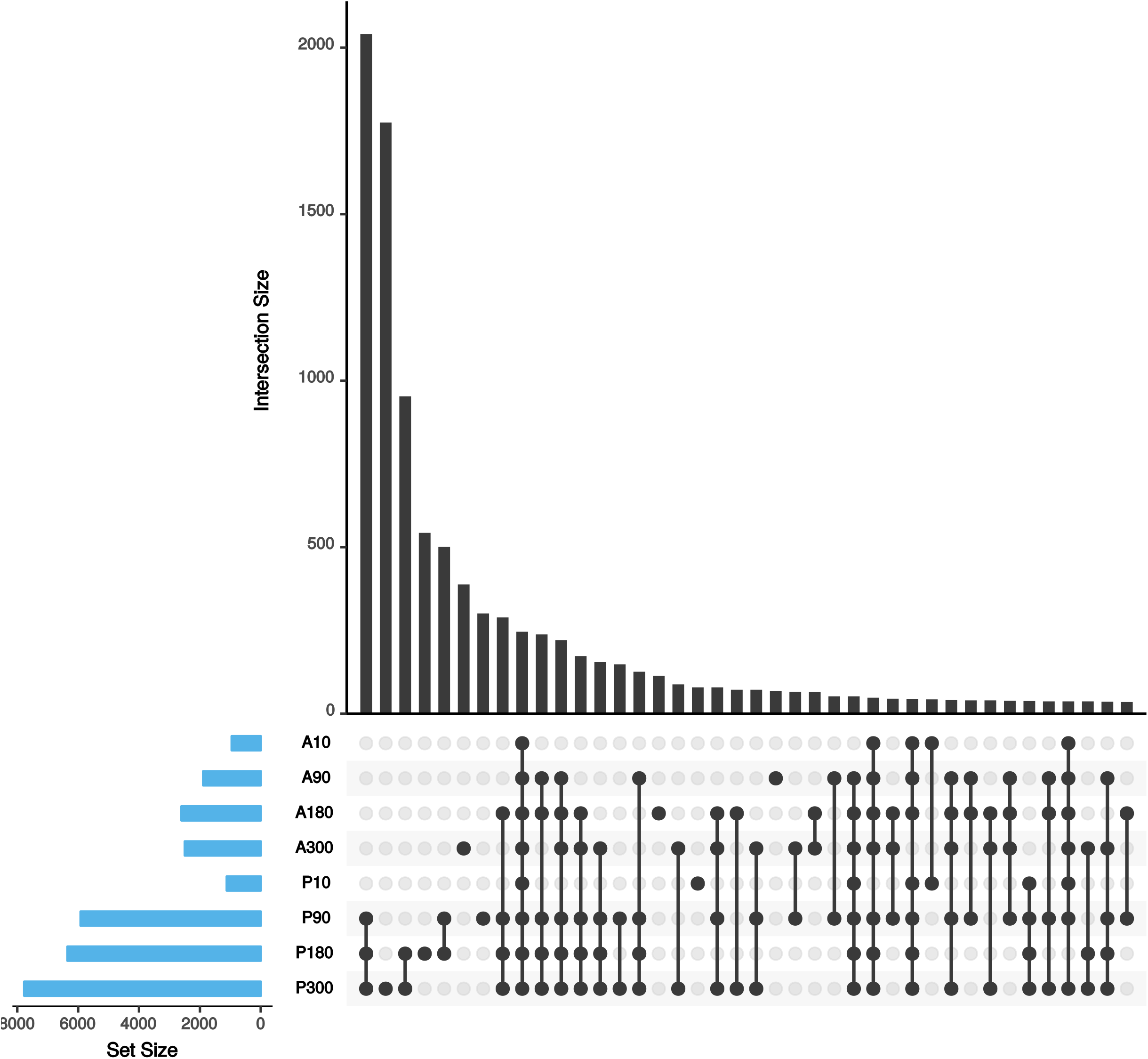
Intersection size between all conditions. The highest number of upregulated genes was found in P300, followed by P180 and P90. The most common regulation pattern for differentially expressed genes was to be upregulated at P90, P180 and P300. See also Figure S1, Table S3 and Table S4.

Judging by the similarity of the expression profiles of regenerating anterior and posterior fragments at 10 minutes post split (A10 and P10; see Fig. 1c), we wondered whether *Stentor* ‘knows’ whether it needs to regenerate an oral apparatus or a hold fast at this stage. We observed that the fraction of shared upregulated genes between A10 and P10 was at least twice as high as compared to upregulated genes of regenerating anterior and posterior fragments for corresponding time points (see Suppl. Information). To analyze this in more detail, all replicates were hierarchically clustered based on their expression patterns (Fig. 3). This analysis revealed that unlike other time points, the A10 and P10 replicates did not form separate clusters (Fig. 3), indicating that similar processes are ongoing at 10 minutes post cell split.

**Figure 3.**
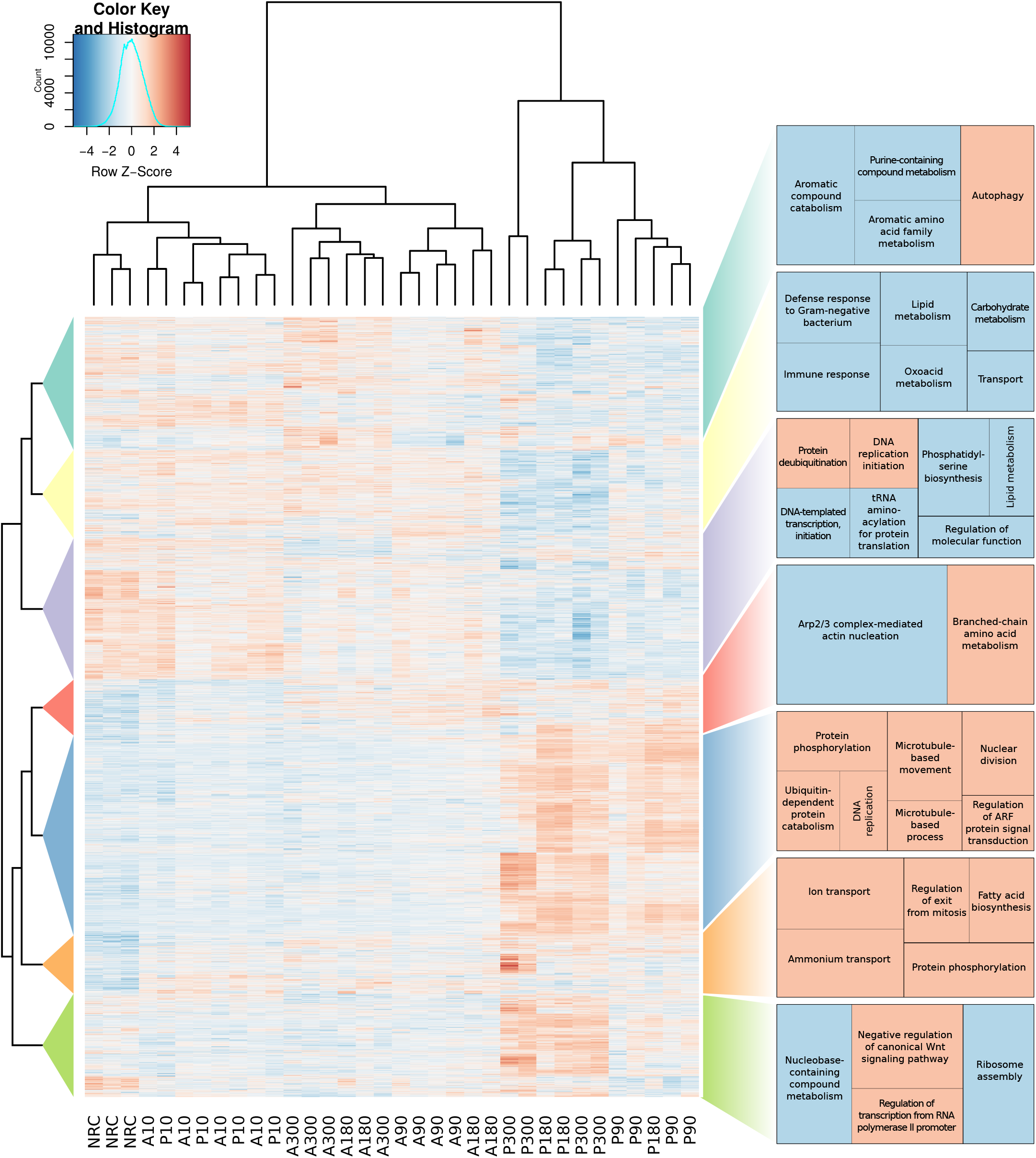
Heatmap with conditions and genes hierarchically clustered. Clustering of both genes and conditions is based on expression patterns. On the heatmap, red indicates high and blue indicate low expression level of the genes. The significantly enriched biological processes for each gene cluster are shown in the side panel. Red and blue background is given to up and down regulated processes, respectively. When *Stentor* is wounded the concentration of ions in the cell will change due to damage in the cell membrane. After *Stentor* has sealed the wound it is expected that ion transport will be needed to re-establish the ion concentration gradients over the membrane. This can also be seen in the heatmap where the cluster with enriched ion transport contains a lot of genes with higher expression levels at 10 min compared to the control. NRC = Non-regenerating cells, A = anterior, P = posterior. See also Figure S2, Table S1, Dataset S1, Dataset S2 and Dataset S3.

A Gene Ontology (GO) enrichment analysis (see Methods for details) was performed to identify any process specific to the anterior or posterior fragment at 10 minutes post split. The analysis showed a high similarity of the processes upregulated in each fragment. The most significant difference between the anterior and posterior fragments were five upregulated proteins involved in cilium movement (GO:0003341), found in the A10 replicates, while only one could be found in P10. Potentially *Stentor* will have less need of such proteins in the P10 condition since the cilia around the oral apparatus are no longer present after the microsurgery. Yet, despite these minor differences, the observed overlap of the A10 and P10 clusters (Fig. 1c) and the shared upregulated processes under these conditions (Fig. 3) indicate that *Stentor* does not yet know what part of the cell it is missing 10 minutes after wounding.

In addition, biological processes specifically upregulated in A10 and P10, that were not present at the later time points were investigated. Processes found to be upregulated at 10 minutes post cell split (phosphorylation, establishment of organelle localization, phospholipid transport, microtubule based movement, potassium ion transport and chitin biosynthesis) could also be found at later time points post cell split. Neither could any downregulated processes specific to A10 and P10 be identified.

The hierarchical clustering of gene expression patterns was then used to further identify functional patterns of regenerating *S. polymorphus* cells (Fig. 3). This approach revealed seven main clusters, four of which were largely dominated by genes that were upregulated for the P90, P180 and P300 conditions; in the three other clusters this pattern was predominantly inverted (Fig. 3). A GO enrichment analysis revealed that each of the clusters was mostly enriched in different biological processes, indicating that genes involved in similar biological processes tend to display similar expression profiles (Fig. 3).

Another GO term enrichment analysis was performed to identify biological processes that were overrepresented in various stages of regenerating *S. polymorphus* cell fragments. Generally, we observe that processes such as signaling, microtubule-based movement, replication, DNA repair and cell cycle as general important processes during cellular regeneration, as these are functionally enriched in many stages of regenerating *Stentor* cells. Conversely, cellular metabolism and processes related to translation, biogenesis and assembly of ribosomes were found to be downregulated under all stages of regenerating cell fragments. This finding indicates that upon cell splitting over the anterior-posterior axis, a functional shift takes place during which typical cellular household functions are downregulated and genes involved in regeneration are activated.

More specifically, we found that three biological processes (GO:0006468 protein phosphorylation, GO:0007018 microtubule-based movement, GO:0006464 cellular protein modification process) were significantly enriched in all stages of regenerating posterior and anterior cell fragments. An orthology-based functional assessment was performed to get more information about the proteins involved in these three biological processes (see Methods for details). The analysis revealed that 16% of the proteins involved in protein phosphorylation were calcium-dependent protein kinases (01IB6, KOG0032, 01KRG), making this the most commonly upregulated kinase family. The proteins assigned as involved in microtubule-based movement were mainly represented by different kinesins (70%) and dyneins (25%). Cellular protein modification process was dominated by tubulin tyrosine ligases (98%) (see Suppl. Dataset 1).

Furthermore, we found that the later stages of regenerating posterior fragments (P90, P180 and P300) shared 20 different enriched biological processes, ten of which being involved in cell cycle related functions (see Suppl. Dataset 2). Similarly, of the five biological processes enriched in the later stages of regenerating anterior fragments (A90, A180 and A300), two were involved in cell cycle progression. These observations indicate that proteins otherwise involved in cell cycle progression are also important for cellular regeneration.

Among the 29 biological functions that were enriched in both middle stages of regenerating posterior fragments (P90 and P180) (see Suppl. Dataset 2), seven were involved in DNA replication or repair, far more than for any of the other stages of regenerating cell fragments. Several of these upregulated genes are involved in DNA synthesis, suggesting that DNA replication takes place during posterior fragment regeneration, but not in regenerating anterior cell fragments. This observation would be compatible with an early pioneering study of regenerating *Stentor* cells, which reported that mitosis took place in regenerating posterior fragments, but not in anterior fragments [4].

Next we used orthology assignments to infer the function of all upregulated genes during various stages of cellular regeneration. This analysis confirmed that genes involved in cell cycle progression were mostly upregulated during the later stages of regeneration of posterior cell fragments (P90, P180 and P300; see Suppl. Fig 1A), but not in other stages of regeneration. Similar trends were observed for genes involved in cytoskeleton remodeling and cell signaling (see Suppl. Fig 1B,C). When only taking the genes assigned to cell cycle into consideration, six of the ten most upregulated genes (based on the average fold change of expression for all conditions; see Methods for details) represented protein kinases. Three of these kinases (SP10077.0, SP11492.0, SP18727.0) were orthologs of the *Saccharomyces cerevisiae* checkpoint kinase SWE1 (KOG0601), which has been shown to act as a negative regulator of entry into mitosis from the G2 phase [6]. The other three kinases (SP17733.0, SP17733.1, SP17733.2) were cyclin dependent kinases belonging to the orthologous group (12T1Q) that includes various kinases that are required for the G1-S transition in mammalian cell cycle [7]. The subunits of Condensin I were all highly upregulated compared to the other cell cycle genes. This complex can induce positive supercoils to DNA during mitotic chromosome condensation [8]. Releasing straining on the DNA helix is probably important for P90, P180 and P300, since all Condensin I subunits were upregulated under these conditions. The CND2 subunit (SP26383.0) was the highest upregulated gene among the cell cycle proteins, and all but one of the remaining subunits in the complex were among the 20 most upregulated when taking all conditions into consideration.

Besides the highly upregulated genes, the orthologous groups that contain the highest number of upregulated genes could potentially represent key regulators of cellular regeneration. We identified 23 orthologous groups that 30 upregulated proteins or more were assigned to (See Suppl. Table 1), among them 9 were groups corresponding to different protein kinases, which is in agreement with the observed enrichment of protein phosphorylation in the GO enrichment analysis.

A total of 103 upregulated genes were assigned as dual-specificity tyrosine-(Y)-phosphorylation regulated kinase (DYRK), making this the group of kinases with the highest number of genes upregulated. The DYRK genes in this orthologous group (KOG0667) were mainly upregulated at P90, P180 and P300. Proteins from both *Tetrahymena* and *Paramecium* were present in KOG0667, however only 7 and 26 paralogues were assigned in these two ciliates, respectively. Additionally no other species in KOG0667 had more than 28 DYRK paralogues, indicating that this large expansion might be a recent event in the evolution of the *Stentor* genus. DYRK is reported to regulate functions such as cell proliferation [9] and phosphorylation of microtubule organizing proteins [10], which are both processes that are highly enriched among upregulated genes in our study. The regulation of these enriched processes together with the wide expansion of the DYRK group indicates the importance of this kinase family during *Stentor* regeneration. A high number of upregulated DYRK genes were also observed in a recent study on regeneration of the oral apparatus in *Stentor coeruleus* [11].

The orthologous group with the highest number of upregulated proteins (0IP8V; see Suppl. Table 1) contains proteins which functions are only poorly understood. These proteins, which were mainly upregulated during P90, P180 and P300, were found to have a high number of MORN domains, and we therefore will refer to this orthologous group as ‘MMDCP’ (Multiple MORN domain containing proteins). MORN domain containing proteins can be found among all major eukaryotic clades, and have been massively expanded among ciliates [12]. The proteins in MMDCP have several MORN domains repeated in tandem, which has been suggested to provide membrane binding capacities [12]. A homology search could identify similarities between MMDCP and the human protein RSPH1, which is another MORN containing protein. Besides the MORN domains, RSPH1 shares the same two domains on the N-terminal end. RSPH1 is suggested to play an important role during meiosis, due to the stage-specific expression of the protein [13]. Interestingly, several MMDCP are upregulated during conjugation in *Tetrahymena* [12], raising the question whether not only proteins active during mitosis, but also proteins involved in meiosis could be implicated in cellular regeneration in *Stentor*. We could identify eight homologues known to have important functions during meiosis that was upregulated during regeneration in *S. polymorphus* (See Suppl. Dataset 3). Three of those, MND1, HOP2 and DMC1 are known to form a meiosis specific protein complex that is involved in double strand break repair. MND1 and HOP2 bind as a heterodimer to DMC1, which is bound to single stranded DNA [14]. Altogether, these observations suggest that several proteins implicated in meiosis are used by *Stentor* during cellular regeneration.

Many of the *Stentor* genes that were upregulated during cellular regeneration lacked a proper annotation due to the lack of identifiable protein homology or protein domains. One of these genes, SP25280.0, was the highest upregulated gene at each of the P90, P180 and P300 conditions. Yet, homology searches of the SP25280.0 protein revealed that it only shared low sequence identity with other protein in the NCBI nr database: the best blast hit for this 1670 amino acid long protein was to a *S. coeruleus* protein (SteCoe_905), which had near full-length coverage (98%), but only relatively low identity (33%). Homologs could be identified in the ciliates *Stylonychia lemnae* and *Oxytricha trifallax*, which both had high query coverage in the homology search. The blastp homology search also found similarities between SP25280.0 and chromosome segregating ATPases based on the identification of an Smc domain on the N-terminal. The highly upregulated SP25280.0 gene, which no function could be inferred for, represents an interesting candidate for future functional studies in *Stentor*.

For regenerating anterior fragments, Lin-54 family protein SP1716.1 was highest upregulated in both the A180 and A300 condition. LIN54 is a subunit of the LINC complex, which regulates genes in cell cycle progression [15]. SP1716.1 shares the Tesmin/TS01-like CXC domain with LIN54, which in the LINC complex is a DNA-binding domain that binds promoters important for cell cycle progression [15]. However, SP1716.1 was found to lack some protein domains that are found in LIN54, and homologues for the other subunits of the LINC complex could not be identified. Therefore, SP1716.1 is unlikely to interact with a complex homologous to LINC, but it is still possible that it is DNA binding like LIN54.

SP23656.0 was the most upregulated gene in P180 among the genes that were not detected as upregulated in any anterior condition. This gene was as well among the most upregulated in P90 and P300. Neither of the two annotation pipelines used could provide an annotation. A blast search against NCBI nr database only gave a hit to *S. coeruleus*, and the same Smc domain as identified for SP25280.0 could also be found. SP1203.1 was among the highest upregulated gene in P90 and P180. At P300 it was the most upregulated gene that at the same was not upregulated at any anterior condition. SP1203.1 got consistently annotated as a transcription factor by both of the annotation software used since it contained the DNA binding E2F/DP family and transcription factor DP domain. These domains are seen together in other transcription factors that regulate cell cycle progression [16]. Therefore SP1203.1 could potentially be a transcriptional regulator of genes involved in regeneration of posterior fragments. Potentially the highest upregulated gene at A90 could be a transcription factor as well. However, for this gene the assumption is only based on the presence of a C2H2 zinc finger domain, which is a nucleic acid binding domain often found in transcription factors [17]. A gene coding for Acetyl-CoA synthetase was detected as upregulated at A180. Interestingly, a phylogenetic analysis containing the most similar bacterial and eukaryotic proteins found in the NCBI database indicated that Stentor acquired this gene via horizontal gene transfer (See Suppl. Fig 2).

Ccr4-Not is a multifunctional protein complex involved in gene regulation by different actions such as mRNA poly(A) deadenylation, tagging of proteins for destruction and acting as a transcription factor [18]. Homologues for all nine subunits found in the *Drosophila* Ccr4-Not were expressed by *S. polymorphus*, and seven of these subunits were found to be upregulated. Surprisingly, the ubiquitin–protein ligase subunit Not4 was shorter in *S. polymorphous*, around half the length compared to homologues in most species. This difference in length were due to the Not4 in *Stentor* was missing a C-terminal domain, which was previously shown by Albert *et al*. [19] to interact with the rest of the Ccr4-Not complex. The Not4 annotation could further be supported by a phylogenetic analysis, which placed the *Stentor* Not4 sequences together with other ciliate Not4 (Suppl. Fig. 3). Almost all of the Ccr4-Not subunits identified were upregulated during P90, P180 and P300, indicating important function in regeneration for posterior fragments at these time points. However, all of the five differentially expressed Not4 homologs were upregulated at other time points, and were following a different expression pattern compared to the other subunits (Fig. 4). This raises the question whether the Not4 homologue in *Stentor* is associated with the Ccr4-Not complex. Taking both the missing complex-binding domain and different expression pattern compared to other subunits into consideration, Not4 in *Stentor* has probably lost the association with the Ccr4-Not. The possibility that Not4 can lose the association with the complex is further supported by an earlier study, which shows that Not4 in human cells are not stably interacting with Ccr4-Not [20]. The potential dissociation of the ubiquitin ligase Not4 indicates a specialization of the Ccr4-Not complex towards gene regulation and degradation of mRNA in *Stentor*.

**Figure 4.**
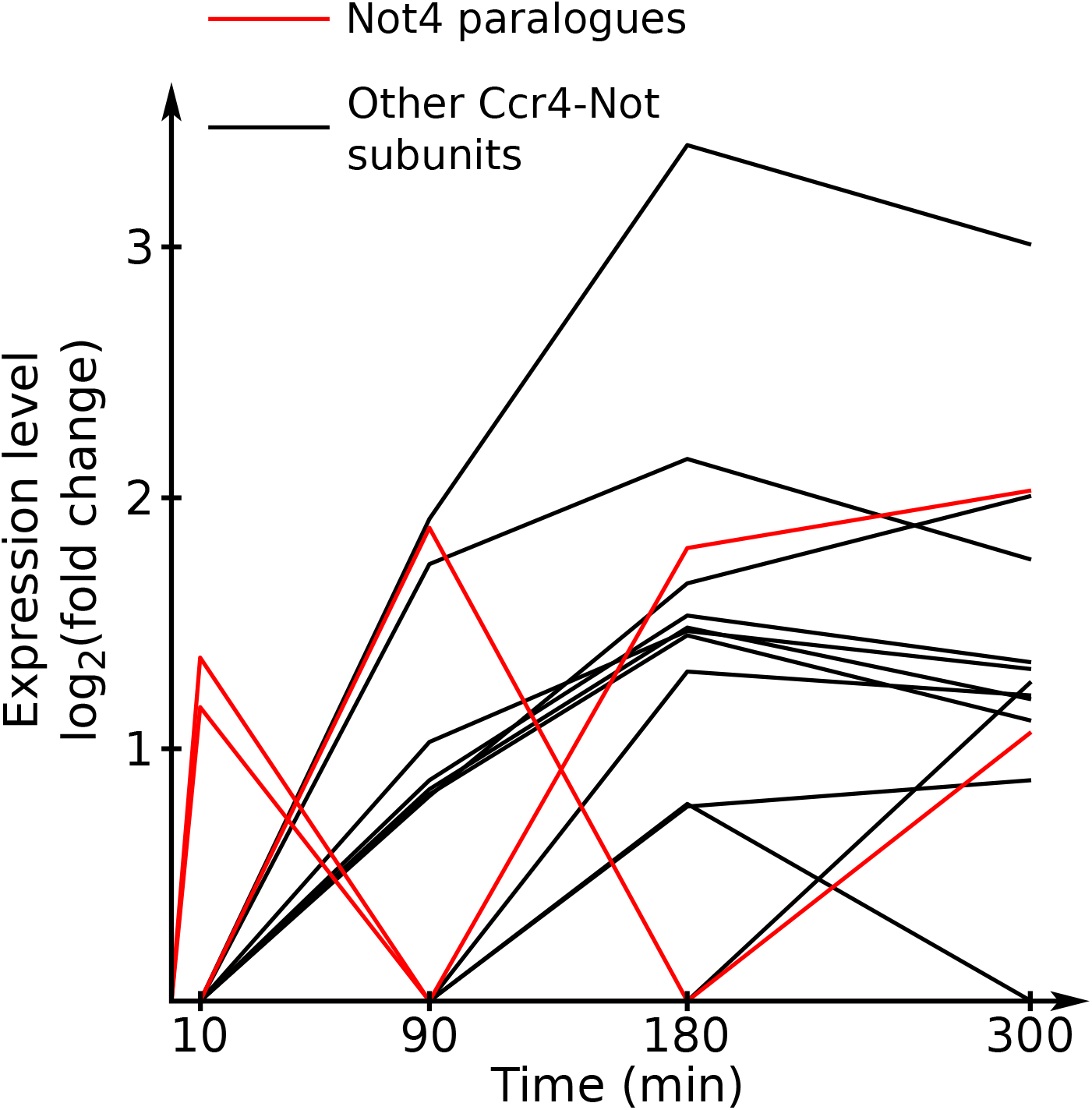
Expression patterns for Ccr4-Not subunits. The five Not4 paralogues, marked with red line, deviate from the expression pattern from the other subunits. See also Figure S3.

Some of the expanded kinesin groups that could be found upregulated in *S. polymorphus* such as DYRK, Aurora kinases and Polo like kinases, are known for their involvement in the cell cycle [9, 21, 22]. In addition, complexes such as condensin and Ccr4-Not indicate that the cell cycle machinery is activated during regeneration. The GO term enrichment study also indicated several cell cycle related biological processes to be enriched. Further investigation of the gene annotation could confirm that homologues known to be important in all phases of the cell cycle, from G1 to telophase, were upregulated (Suppl. Dataset 3). This confirms that regeneration have many similarities to cell division, which has earlier been observed in early morphological and cell biological studies of regenerating *Stentor* cells [3, 4]. Based on the upregulated expression of MORN domain containing proteins, the meiosis specific MND1, HOP2, DMC1 and other proteins involved in the cell cycle (Suppl. Dataset 3), regeneration also share some similarities with meiosis.

With *Stentor* being a model organism for studying cell biological and molecular processes involved in cellular regeneration, the recent release of the *S. coeruleus* genome represented an important milestone [5]. In the current study we have used an RNAseq approach to identify genes that are involved in the regeneration of *S. polymorphus* cells. These genes, and especially those that were highly upregulated during cellular regeneration but lack a proper functional annotation, represent interesting candidates for further studies that aim to elucidate key factors in this process. This study also shows that transcriptome data can be generated from *Stentor* cell fragments, opening up avenues for future studies that aim at investigating transcriptional responses below the level of individual cells.

## Author Contributions

H. O.G. and T.J.G.E. conceived the study. H.O.G. and M.J. performed sampling, laboratory and bioinformatics analyses. H.O.G. and T.J.G.E. wrote the manuscript and all authors read, edited and approved the final manuscript.

## Acknowledgements

All sequencing was performed by the National Genomics Infrastructure sequencing platforms at the Science for Life Laboratory at Uppsala University, a national infrastructure supported by the Swedish Research Council (VR-RFI) and the Knut and Alice Wallenberg Foundation. This work was supported by grants of the Swedish Research Council (VR grant 621-2009-4813), the European Research Council (ERC Starting grant 310039-PUZZLE_CELL) and the Swedish Foundation for Strategic Research (SSF-FFL5) to T.J.G.E.

## Method details

### Sample collection and generation of sequencing data

Sediment samples were collected with a 500 ml cage sampler from a pond in Uppsala (59^o^50’19.0”N 17^o^37’21.4”E). The bottles were taken back to the lab within 30 minutes. When the debris had settled, sub samples from the bottles were taken with a transfer pipette from the interface between sediment and water and moved into a petri dish. Under a stereo microscope *Stentor polymorphus* could be identified and single cells could be moved into another petri dish containing 2% methylcellulose solution. Using a scalpel *S. polymorphus* cells were divided in the middle, separating the anterior (part of *S. polymorphus* containing the oral apparatus) from the posterior (part of *S. polymorphus* containing the hold fast). Cell fragments were moved into 15 ml centrifuge tubes containing pond water passed through a 5 μm filter. The tubes were degassed with nitrogen, tightly sealed and stored in dark until they were picked in a 1 μm volume and put in a petri dish containing sterile water. Single cells were then picked in a 0.5 μl volume into sterile 0.2 ml PCR tubes. After picking each cell, this cell was immediately stored in -80 ^o^C. This procedure was done multiple times using different incubation times for the cells in the filtered pond water, i.e. varying the regeneration time prior to -80 ^o^C storage. The different times for incubation in degassed tubes were 90 min, 3 h and 5 h. Replicates with 10 min incubation time post microsurgery were prepared in the same way with the exception that cells were never transferred into degassed tubes, but were kept in a petri dish with filtered pond water until the washing step followed by -80 ^o^C storage. Four replicates of both anterior and posterior fragments at the different time points were prepared. As controls, four replicates of whole *S. polymorphus*, excluding the microsurgery step were picked and 3 replicates of dividing cells at the stage referred to as “step 7” by Tartar [23]. In total 39 replicates were prepared, for these tubes containing single cells or single cell fragments cDNA were generated and prepared for sequencing according to the Smart-seq2 protocol [24]. The Nextera XT libraries were sequenced on a HiSeq2500 (Illumina), using rapid mode setting, generating paired end 250 bp reads. Among the 39 cDNA libraries prepared, 112 Gb sequencing data was successfully produced for 34 of the replicates included in the final workflow. Leading to only three replicates for non regenerating cells and three replicates of P180 that were analyzed together with four of each of the other conditions. The dividing replicates where picked from a petri dish with a high density of *S. polymorphus* monitored for approximately 60 minutes under light, therefore these replicates also spent longer time in high oxygen conditions than the other replicates. The dividing cells were also excluded since the differences in experimental background compared to the other replicates made the downstream data analysis challenging.

### Amplicon sequencing with universal 18S rRNA primers

DNA was extracted from 0.25 g of bottom sediments from the Stentor sample site using the PowerSoil extraction kit (MoBio). The extracted DNA prepared for sequencing according to a method described earlier by Hugerth et al. [25], using the 18S directed primer pair 574*f and 1132r. The sequencing of the 18S rRNA amplicons was performed on a MiSeq sequencer (Illumina), generating paired end 250 bp reads. Because of anticipated polymerase and sequencing errors, clustering of OTUs (Operational Taxonomic Units) was done with a 97% threshold [26]. Sequencing of the amplicons revealed a single dominant *Stentor* cluster containing more than 99% of reads assigned to the *Stentor* genus. All Stentor 18S rRNA gene sequences found in the transcriptomes generated were assigned to this *Stentor* OTU.

## Quantification and statistical analysis

### *De novo* transcriptome assembly and differential expression analysis

All raw sequencing reads were assembled into a co-assembly using Trinity v2.2.0 with the *in silico* normalization of reads option [27]. Smart-seq2 primers and Nextera XT adaptors were removed with the Trimmomatic version bundled with Trinity applying the settings “ILLUMINACLIP:2:30:10”. Also in the following order LEADING:5, TRAILING:5 SLIDINGWINDOW:5:16 and MINLEN:80 was applied to trim low sequencing quality regions from reads. To map back the reads to the coassembly Bowtie2 v2.2.3 [28] with the settings “--end-to-end -k 20 -D 20 -R 3 -N 1 - L 20 -i S,1,0.50 -p 16 -X 1000” was used. Genes were clustered and a count matrix for the differential expression analysis was made with Corset [29] dividing the data into the 9 different experimental groups. In the corset run links between contigs were filtered out if the link was supported by less than 3 reads, all other settings were default. Differentially expressed genes in each condition compared to the control were identified with DeSeq2 [30]. In a previous study, a kinase named mob1 has been reported as a regulator of regeneration [31]. Our differential expression analysis detected mob1 as upregulated in both anterior and posterior fragments at 90, 180 and 300 min post split of cell, which is consistent with earlier observations. To rank which gene was most upregulated while taking all conditions into consideration the sum of all fold changes for each gene were sorted in descending order.

### Assessment of potential contaminant genes

Potential contamination levels were assessed by a diamond aligner blastx search [32] against NCBI nr database. The longest contig from each corset cluster was used as query. When blasting the whole co-assembly against nr, 93% of the genes had a top hit to *Stentor coeruleus* (Table S2). The organism that had the second most top blast hits was *Chlorella variabilis*. This is not surprising, since *S. polymorphus* contain green algal endosymbionts. When blasting all the genes assigned as differentially expressed, 96% of these genes had their top hit to *S. coeruleus* (Table S3). The five species with most top hits in the diamond alignment to nr were all ciliates. This indicates that the data used in the differential expression analysis contained a low level of contamination.

### Annotation and GO enrichment analysis

The co-assembly was translated in all possible reading frames using TAA and TAG as stop codons. We did not use TGA as a stop codon since it could not be found in the initial 160 genes that were investigated to assess codon usage. After the discovery that *Stentor* do use TGA as stop codon [5], we still kept our ORF assignment since the TGA stop codon was so rare and read-through often lead to other stop codons in tandem after the first one or the assembled transcript ended anyways. The longest open reading frame on each contig and each open reading frame longer than 80 amino acids were annotated with the first public beta release of eggNOG-mapper [33] and InterProScan v5.23-62.0 [34] using the databases SUPERFAMILY, Hamap, ProSiteProfiles, TIGRFAM, SMART, CDD, PRINTS, PIRSF, ProSitePatterns, Pfam and PANTHER. The GO terms assigned by InterProScan were used to perform a GO enrichment analysis with topGO developed by Alexa & Rahnenfuhrer in 2016. The annotations of the upregulated proteins were curated manually by inspection of mafft-linsi alignments [35] with the annotated proteins from *S. polymorphous* and proteins from the orthologous groups assigned by eggNOG-mapper [33, 36]. Annotations where alignment was poor or major conserved domains were missing were discarded. To support that *Stentor* express a high number of different genes from the reported orthologous group, these genes were blasted against the *S. coeruleus* genome. The blast search could then confirm the presence of similar number of genes in the *S. coeruleus* genome as observed upregulated in *S. polymorphus*. Only curated annotations were used throughout the study, except for when the highest upregulated genes where investigated without taking function into consideration.

### Phylogenetic analyses

The *S. coeruleus* homologues were identified via using the *S. polymorphus* sequences as blast queries against the *S. coeruleus* proteome [5]. In the Not4 phylogeny proteins from the orthologous group KOG2068 was used in the multiple sequence alignment. For the Acetyl-CoA phylogeny the top 250 eukaryotic and bacterial blasthits against NCBI nr database were used. The alignments were curated from sequences representing long branches that potentially could cause artefacts in the tree topology. The curated alignment was trimmed with trimAl v1.4.rev15 [37] using 0.4 as gap threshold for the Acetyl-Coa phylogeny and 0.5 for Not4. The tree topologies were calculated using ultrafast bootstraps with IQ-tree v1.5.3 [38]. The models were selected using the bayesian information criterion.

